# Sex differences, asymmetry and age-related white matter development in infants and 5-year-olds as assessed with Tract-Based Spatial Statistics

**DOI:** 10.1101/2023.01.29.526138

**Authors:** Venla Kumpulainen, Harri Merisaari, Eero Silver, Anni Copeland, Elmo P. Pulli, John D Lewis, Ekaterina Saukko, Satu J. Shulist, Jani Saunavaara, Riitta Parkkola, Tuire Lähdesmäki, Linnea Karlsson, Hasse Karlsson, Jetro J. Tuulari

**Author notes:** shared contribution (Eero Silver, Anni Copeland, Elmo P. Pulli). Corresponding author, Venla Kumpulainen, Kiinamyllynkatu 10, FinnBrain study, Medisiina A Building, 20520 Turku.

## Abstract

The rapid white matter (WM) maturation of first years of life is followed by slower yet long-lasting development, accompanied by learning of more elaborate skills. By the age of five years, behavioural and cognitive differences between females and males, and functions associated with brain lateralization such as language skills are appearing. Diffusion tensor imaging (DTI) can be used to quantify fractional anisotropy (FA) within the WM and increasing values correspond to advancing brain development. To investigate the normal features of WM development during early childhood, we gathered a DTI data set of 166 healthy infants (mean 3.8 wk, range 2-5wk; 89 males; born on gestational week 36 or later) and 144 healthy children (mean 5.4 years, range 5.1-5.8 years; 76 males). The sex differences, lateralization patterns and age-dependent changes were examined using tract-based spatial statistics (TBSS). In 5-year-olds, females showed higher FA in wide-spread regions in the posterior and the temporal WM and more so in the right hemisphere, while sex differences were not detected in infants. Gestational age showed stronger association with FA values compared to age after birth in infants. Additionally, child age at scan associated positively with FA around the age of 5 years in the body of corpus callosum, the connections of which are important especially for sensory and motor functions. Lastly, asymmetry of WM microstructure was detected already in infants, yet significant changes in lateralization pattern seems to occur during early childhood, and in 5-year-olds the pattern already resembles adult-like WM asymmetry.

**Highlights:**

- White matter tract integrity shows widespread sex differences at the age of 5 years.
- White matter structure is highly lateralized during early childhood, and changes in asymmetry occur between the birth and 5 years of age.
- The white matter lateralization pattern of 5-year-olds, unlike of infants, resembles asymmetry observed in adults.

## 1 Introduction

White matter (WM) microstructural development is linear during prenatal period^1^ and continues throughout the childhood with widespread increase in fractional anisotropy (FA) and corresponding decrease in mean diffusivity (MD)^2–14^. The remodelling follows specific trajectories with both regional and temporal variation reflecting the concomitant maturation of motor, behavioural, and cognitive skills.

Generally, postnatal brain development advances with posterior-to-anterior and central-to-peripheral, non-linear patterns^2^ and continues from the second trimester into adulthood. Especially rapid developmental period occurs during the first two years of life, when myelination and pruning of axonal connections are driving the increase in WM integrity ^3,15,16^. The myelination mainly limits the diffusion perpendicular to the axon axis, which is detected as decreasing radial diffusivity^17^ and concomitantly increasing FA values during the development ^2–5,8–12,16,18–31^. Commissural and projection fibres, including corpus callosum and pyramidal tracts, are first to reach the adult-like integrity before school-age ^10,12,26,32^. These tracts are related to improvement of both gross and fine motors skills, which are prerequisites for further learning of more elaborate skills like writing ^33,34 35^. Subsequently, the connections from occipital and limbic areas, associated with e.g. the development of language skills, mature between ages five to eight years ^36–38^, followed by the maturation of fronto-temporal association tracts, related to higher-order functions ^39–44^, with prolonged development continuing to early adulthood ^10^. However, the exact timing of changes especially in young children is still partly elusive and warrants further study.

The developmental patterns of WM show sexual dimorphism with earlier maturation detected in females ^32,44–47^, yet more rapid and longer-lasting age-dependent increases in FA in males contributes to levelling off the differences by the age between 10 and 14 years^47^. The studies have provided varying results on the divergence of the trajectories, both in the timing and tract-specific features (Table 1). In two studies, sex differences were already detected during neonatal age^48,49^, even though the differences were limited to few areas. In a sample of 25 infants, males had lower FA in right corticospinal tract^48^, and in another study of age range 0 to 2 years^28^, females showed higher FA in the right sensory and the right uncinate tracts and lower FA in the arcuate and motor tracts. Differences in WM development between females and males were detected in multiple study populations of older children and adolescents ^21,32,45,46,50^, and more specifically between the ages of two and eight years ^10^. On the other hand, no developmental sex differences were detected in some studies with wide age ranges ^14,26,51,52^. Furthermore, part of the studies have shown FA values in specific tracts (e.g. in the corticospinal tract (CST) ^10,21,32,46^) to be higher in males during childhood, in contrast to generally lower FA values detected in males. Additionally, the lateralization of WM microstructure is well-acknowledged both during the brain development ^19,29,30,51,53^ and adulthood ^37,54–57^. Asymmetry of WM tract diffusion properties has been shown to emerge as early as during neonatal age ^58^, and is associated with later development of, e.g., language skills ^59^ but is a relatively understudied topic.

**Table 1.**
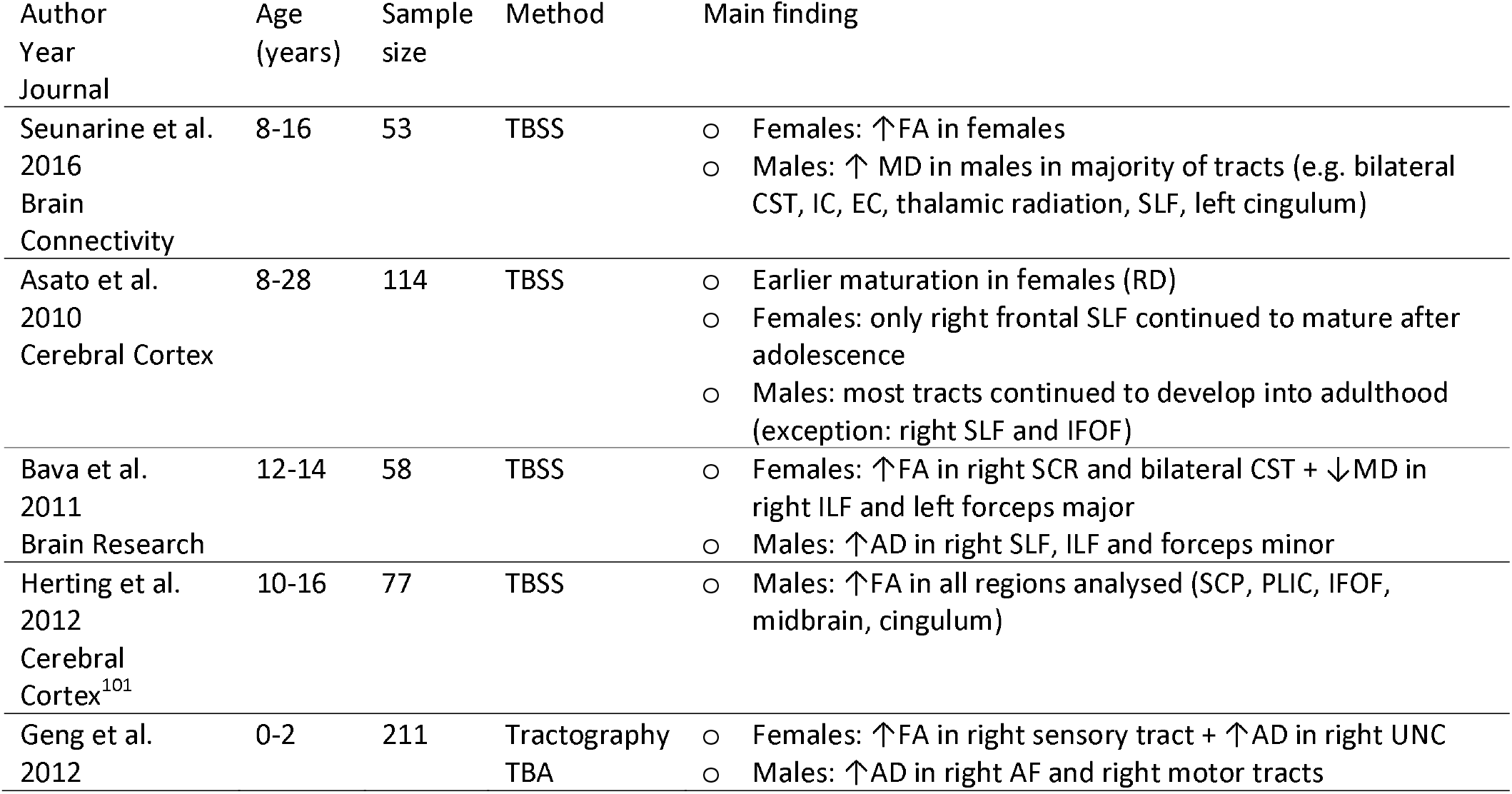
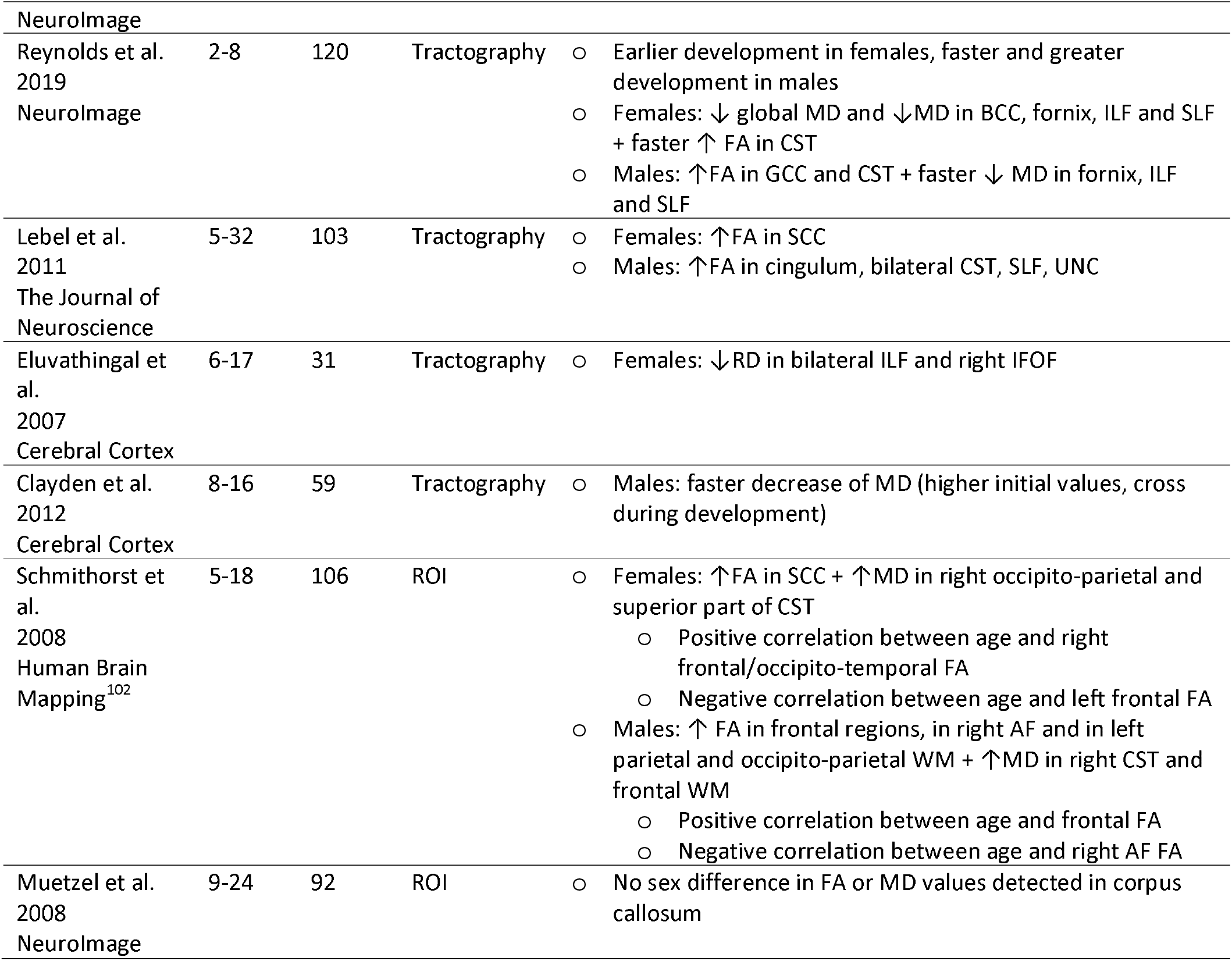
Literature review of white matter sex differences in age groups close to our study. TBSS = tract-based spatial statistics, TBA = tract-based analysis, ROI = region of interest, FA = fractional anisotropy, MD = mean diffusivity, RD = radial diffusivity, AD = axial diffusivity, CST = corticospinal tract, IC = internal capsule, SLF = superior longitudinal fasciculus, IFOF = inferior fronto-occipital fasciculus, ILF = inferior longitudinal fasciculus, SCP = superior cerebral peduncle, PLIC = posterior limb of internal capsule, UNC = uncinate fasciculus, AF = arcuate fasciculus, BCC = body of corpus callosum, GCC = genu of corpus callosum, SCC = splenium of corpus callosum

Sex differences in emotional processing^60,61^, personality features^62,63^, temperament^64–66^ and behaviour^67,68^ are widely reported during childhood and adolescence. Additionally, prevalence of certain diseases is related to sex, e.g., females show more internalizing symptoms and associated disorders such as anxiety^69,70^ while incidence of antisocial behaviour^71^ and diagnosis of attention deficit hyperactivity disorder (ADHD)^72,73^ are overrepresented in males. Unravelling sex-associated features of brain structure and function provide important knowledge on the factors inducing these sex-specific cognitive patterns. Furthermore, alterations in WM lateralization have been associated with conditions such as autism^74^ and ADHD^75^. Distinguishing normal variability of WM lateralization might open up new venues for characterizing also disease-associated changes.

Here, we describe sex differences, lateralization patterns, and developmental changes of the brain WM microstructure in populations of infants and 5-year-olds. We hypothesized to find higher FA values in females, positive association between FA values and age, and leftward asymmetry in both age groups based on prior literature.

## 2 Materials and methods

This study was reviewed and approved by the Ethics Committee of the Hospital District of Southwest Finland ((07.08.2018) §330, ETMK: 31/180/2011), and performed in accordance with the Declaration of Helsinki.

### 2.1 Participants

The study included two subpopulations of families recruited as a part of FinnBrain Birth Cohort^76^ to brain magnetic resonance imaging (MRI). Twenty-two subjects were included in both subpopulations (10 males). The exclusion criteria of both subpopulations were set to screen healthy participants with no acknowledged developmental problems. The first subpopulation of 190 infants (age of 11 to 54 days from birth at scan, mean 27, SD 7.9; 103 males and 87 females) participated in brain MR scanning between December 2012 and January 2016. The exclusion criteria for infants were: 1) Birth before gestational week (gwk) 36, 2) birth weight of less than 2500g, 3) Apgar score < 5 at 5 minutes after birth, 4) previously diagnosed central nervous system anomaly, an abnormal MRI finding in any previous MRI scan or severe perinatal complication with neurological consequences.

The MR scans of 203 5-year-olds were performed between October 2017 and March 2021. The participants were healthy, normally developing children aged from 5.1 to 5.8 years (mean 5.4, SD 0.11, 111 males, 92 females, 182 right-handed, 14 left-handed, 7 with no preference). The exclusion criteria, in addition to general MRI contraindications, were the following: 1) Birth before gwk 35 (before gwk 32 for separate nested cohort with exposure to synthetic glucocorticoids during pregnancy), 2) major developmental disorder, 3) other types of long-term diagnosis that requires constant contact to hospital, 4) sensory abnormalities (e.g. blindness or deafness), 5) use of daily, regular medication (asthma inhalers during infections was not an exclusion criterium; and desmopressin (^®^Minirin) medication was allowed), 6) head trauma requiring inpatient care (reported by parents). The informed consent was collected from both parents.

### 2.2 MR imaging

Data acquisition was conducted in Turku University Hospital with a Siemens Magnetom Verio 3T scanner (12-element head/neck matrix coil) for infants and with a Siemens Magnetom Skyra fit 3T scanner (20-element head/neck matrix coil) for 5-year-olds. The scanner was upgraded in between the two data collections. The Generalized Autocalibrating Partially Parallel Acquisition (GRAPPA) technique was used to accelerate imaging. Diffusion weighted imaging protocol was applied with a standard twice-refocused Spin Echo-Echo Planar Imaging (SE-EPI): TR (time of repetition)/TE (time of echo) = 8500/90.0ms (for infants) and 9300/87.0ms (for 5-year-olds), FOV (field of view) = 208mm, isotropic voxels with 2.0×2.0×2.0mm resolution, b value 1000s/mm^2^, 96 noncollinear diffusion gradient directions divided into three scanning sets (31, 32, and 33 directions) with 9 b0 images (3 b0 = 0s/mm^2^ volumes scattered at the beginning, middle and the end of the set). The scan visits included also structural and functional MRI scans^77^, but the current study uses only the structural diffusion data.

The infants were imaged during natural sleep after being fed by (breast) milk. A vacuum mattress was used to reduce movement during the imaging. Soft mouldable silicone ear plugs and hearing muffs were used for hearing protection. Infant MR images included 12 subjects with incidental findings (mainly small subdural hemorrhages), which were handled according to the procedure described in our earlier work^78^ and were regarded not to affect the results.

The imaging visit of 5-year-olds was preceded by a home visit to provide detailed information on the study and a home practice period. The imaging session was also simulated before scanning, including practices of lying still and learning how to communicate during imaging. All children were imaged awake or during natural sleep. For more detailed description of the whole image set, the practice period and the imaging visit, see the previous published studies ^77,79,80^.

### 2.3 Data analysis

DTI data acquisition was successful with 169 infants and with 163 5-year-olds. We used default settings in DTIprep denoising, slice-wise intensity checking, interlace-wise venetian blind artefact setting, and baseline averaging prior to porting the denoised and cleaned data for tensor fitting with FSL tools (https://nitrc.org/projects/dtiprep/ ^81^). As the applied settings were detected to pass volumes with motion-related artifacts, all volumes were manually inspected and the ones with residual artifacts were excluded. For subsequent analysis, 30 directions were chosen per subject by maximizing the angular resolution, and subjects with insufficient number of directions after quality control steps were excluded ^82^. We chose to use 30 directions, as in our previous study^79^ we observed that increasing the number of diffusion-encoding directions over 18 provided no additional advantage in repeatability of results. Further, inclusion of more directions would have led to a decrease in the sample size. With this choice, the final number of subjects was 166 infants (89 males, 77 females) and 144 5-year-olds (76 males, 68 females).

Pre-processing of quality-controlled data was conducted with FSL 6.0 (FMRIB software library, University of Oxford, UK). b0 images, including at least one image per imaging set, were co-registered and averaged (FLIRT, FMRIB’s Linear Image Registration Tool), and the brain mask was created with Brain Extraction Tool (BET version 1.0.0)^83^ with settings -R -f 0.3. Motion and eddy current correction were performed with FSL tools with concurrent rotation of the b vector matrix. The scalar maps were computed with FSL’s dtifit. Tract-based spatial statistics (TBSS) pipeline of FSL ^84^ was used to estimate WM tract skeletons with the following settings (5-year-olds/infants): “tbss_2_reg” flag -T/-t (target being a study-specific template), “tbss_3_postreg” flag -S, FA threshold 0.2/0.15. Relative residual rotational and translational framewise displacements were measured to quantify the intrascanner head motion. The used pipelines have been thoroughly validated, please see more detailed description of data analysis pipeline, test-retest reliability, and assessment of optimal number of diffusion-encoding directions to DTI scalars in our previously published articles ^79,82,85^.

General linear model (GLM) with FSL’s randomise tool was used for voxel-wise analysis of inter-subject variation. In the infant group, sex, age from birth and gestational age, rotational and translational mean framewise displacement measures (to control intrascanner head motion), maternal pre-pregnancy body mass index (BMI), birth weight, maternal smoking during pregnancy, and maternal socio-economic status (SES; evaluated by educational level and categorized into two classes: 1. low [elementary school] and medium [high school or occupational education], 2. high [examination from university or university for applied sciences]) were explored as independent contributors. For 5-year-olds, sex, age at scan, handedness, rotational and translational mean framewise displacement measures, maternal pre-pregnancy BMI, ponderal index, maternal smoking during pregnancy and maternal SES were explored as independent contributors. The analyses were conducted with 5000 permutations and multiple comparison correction with threshold-free cluster enhancement (TFCE; corrected p < 0.05 regarded as statistically significant). As the age and sex showed statistically significant correlations with FA, further analyses were performed with age and sex as main variables and other variables as covariates.

Sensitivity analyses with gestational age (birth before gestational week 35 for 5-year-olds), maternal pre-pregnancy BMI, maternal age at birth, maternal SES, smoking during pregnancy, prenatal exposure to corticosteroids or selective serotonin/serotonin-noradrenaline reuptake inhibitors (SSRI/SNRI) during pregnancy, and maternal perinatal depressive/anxiety symptoms (Edinburgh Postnatal Depressive Scale (EPDS)^86^ and Symptom Checklist (SCL-90)^87^) at 2^nd^ trimester and three months postpartum (for 5-year-olds) were conducted. See the supplementary material for demographical information and regression analysis workflow (Supplement Tables 1 and 2).

Additionally, we examined the lateralization patterns of FA by FSL’s “tbss_sym” both over all participants and separately in females and males. Lateralization was estimated by subtracting the right-sided FA values from the left-side ones, followed by a “one-sample t test” randomise analysis (5000 permutations, TFCE correction) to detect areas with significant variation. The mean image and standard deviation (SD) of lateralized values was calculated with fslmaths. Tract-wise mean FA values were extracted with coregistration of JHU Neonate atlas^88^ for infants and JHU-ICBM-DTI atlas^89^ for 5-year-olds to the TBSS skeleton images.

Statistical analyses for tract-wise extracted mean FA values were performed with SPSS version 27.0 (IBM Corp. 2020, Armonk, NY, USA). Tract-wise sex differences were estimated with independent samples t test (two-tailed), and asymmetric differences with one-sample t test, with Bonferroni corrected p value thresholded at 0.001 (0.05/36 tracts). Effect sizes for group differences were estimated with Cohen’s d. Comparison between asymmetries in a subcohort of 22 infants and 5-year-olds was conducted with paired t test with Bonferroni corrected p value threshold at 0.002 (0.05/22).

## 3 Results

### 3.1 Sex differences

#### 3.1.1 Infants

Sex showed no statistically significant correlation with FA values in the infants after controlling for age at scan and gestational age, intrascanner head motion, birth weight, maternal BMI, SES, and smoking.

#### 3.1.2 Five-year-olds

A significant effect of sex on FA values was detected in widespread regions at the age of five years, and was statistically strongest in posterior and temporal parts, and in the right hemisphere (Figure 1). Females had higher FA values in all regions with significant difference, i.e. no regions with higher FA in males were detected. The results remained significant after controlling for gestational age, handedness, ponderal index, intrascanner head motion, smoking during pregnancy, pre-pregnancy BMI, and SES. In sensitivity analyses, the pattern of results in TBSS remained similar and statistically significant (data not shown). Group differences of whole WM tract mean FA values between females and males are provided in Supplement Table 3.

**Figure 1.** Sex-differences with tract-based spatial statistics (TBSS) at the age of 5 years. Higher fractional anisotropy (FA) values detected in females are present in widely distributed areas, especially in posterior and right hemisphere regions. Regions with statistically significant difference showed in 3D (top left corner) and in axial plane (bottom left), threshold-free cluster enhancement (TFCE) correction applied, regression with 5000 permutations, p < 0.05 (red colour bar showing p values), age as a covariate. Additionally, on the right, group-differences in mean FA across whole tract between females (F) and males (M) showed in splenium of corpus callosum (SCC; Cohen’s d 0.48), right retrolenticular internal capsule (rl IC; Cohen’s d 0.58), left inferior longitudinal fasciculus/inferior fronto-occipital fasciculus (ILF/IFOF; Cohen’s d 0.58) and left posterior thalamic radiation and optic tract (PTR/OR; Cohen’s d 0.66). p < 0.005 = *, p < 0.001 = **. R = right, L = left

### 3.2 Age effects

#### 3.2.1 Infants

The gestational age showed wide statistically significant positive correlations (p < 0.01) with FA values after controlling for age from birth and sex (Figure 2; results with p < 0.05 in the Supplement Figure 1). Additionally, age from birth showed positive correlations with FA values (p < 0.05) after controlling for the gestational age and sex. The effect of gestational age on FA values was greater compared to the effects detected after birth. The results remain significant after controlling for intrascanner head motion, prenatal exposure to smoking, maternal age, pre-pregnancy BMI and SES. The covariates included in the sensitivity analyses did not affect the results. We inspected the age effects also separately in females and males and observed no difference in association between gestational age and FA after controlling for age after birth or other covariates between males and females. When inspected separately, the age after birth and FA showed positive association in males (p < 0.05 after controlling for covariates), but the result did not remain statistically significant in females.

**Figure 2.** Effect of the age (controlled with gestational age) and the gestational age (controlled with age after birth) on fractional anisotropy (FA) with tract-based spatial statistics (TBSS) in infants. Positive correlation between both ages and FA are detected, with stronger and wider correlation between gestational age and FA. TFCE (threshold-free cluster enhancement) correction applied, 5000 permutations, p < 0.05 (for age from birth) and p < 0.001 (for gestational age), blue colour bar showing p value. R = right, L = left.

#### 3.2.2 Five-year-olds

Within our age range of 5.1 to 5.8 years, the age was positively correlated with FA values in the body of corpus callosum (BCC) (Figure 3). After controlling for maternal pre-pregnancy BMI, maternal age, SES, and exposure to tobacco smoking during pregnancy, the result did not remain significant (TFCE corrected p < 0.06, with which also an additional cluster in left frontal area was observed; See Supplement Figure 2). When examined separately, no significant age-effects were found in males, and in females the effect in BCC did not reach statistical significance (p = 0.08).

**Figure 3.** Effect of the age on fractional anisotropy (FA) with tract-based spatial statistics (TBSS) around the age of 5 years. Positive correlation between age and FA of corpus callosum is detected. TFCE (threshold-free cluster enhancement) correction applied, 5000 permutations, p < 0.05 (red-yellow colour bar showing p value), age as a covariate. P = posterior, A = anterior, R = right, L = left.

### 3.3 Lateralization

#### 3.3.1 Infants

Lateralization of brain WM was detected in infants in multiple tracts. Leftward lateralized WM regions are depicted in Figure 4 (TFCE corrected p < 0.01; results with p < 0.05 in the Supplement Figure 3). Sex was not observed to affect the lateralization pattern.

**Figure 4.** Lateralization pattern of white matter tract fractional anisotropy (FA) with tract-based spatial statistics (TBSS) in infants. Significant regions with leftward asymmetry in blue. Threshold-free cluster enhancement (TFCE) correction applied, 5000 permutations, p < 0.01 (blue colour bar showing p value). R = right, L = left, A = anterior, P = posterior. Tracts are marked with white circles. Additionally, left-right difference (left minus right, negative values denote rightward asymmetry) of FA in uncinate (UNC), anterior limb of internal capsule (ALIC), posterior limb of internal capsule (PLIC), thalamic radiation/optic tract (PTR/OR), posterior and superior corona radiata (PCR and SCR), and the statistical significance with one-sample t test and effect sizes.

#### 3.3.2 Five-year-olds

The WM structure was highly lateralized at the age of five years (Figure 5). Statistically significantly higher FA values in left-sided tracts were detected in multiple areas (e.g. frontal and temporal areas and in midbrain; TFCE corrected p < 0.001 in Figure 5 and with p < 0.05 in the Supplement Figure 4). The mean image of lateralization pattern (Supplement Figures 5 and 6) indicating the magnitude and direction (left/rightward) of the hemispheric asymmetry of FA values also shows rightward lateralized regions, for example, in the anterior limb of internal capsule (ALIC) and antero-lateral corpus callosum. Leftward asymmetry was detected, for example, in cingulate, inferior fronto-occipital fasciculus, posterior corpus callosum, fronto-parietal region, and midbrain. In most of the WM tracts, both leftward and rightward lateralized subregions were observed. Inter-subject variation of FA asymmetry was detected to be emphasized to peripheral WM tract regions (Supplement Figure 7 shows the SD of lateralization pattern). Tract-wise lateralized FA (left-right) and effect sizes for the differences between whole tract mean FA values are provided in Supplement Table 4. Statistically significant differences in the asymmetry pattern were not observed between females and males (Supplement Figure 8 shows sex-specific left-right lateralization patterns).

**Figure 5.** Lateralization pattern of white matter (WM) tract fractional anisotropy (FA) with tract-based spatial statistics (TBSS) around the age of 5 years. Significant regions with leftward asymmetry in red, threshold-free cluster enhancement (TFCE) correction applied, 5000 permutations, p < 0.001 (red colour bar showing p value). R = right, L = left. Specified tracts are marked with white circles. Additionally, left-right difference (left minus right, negative values denote rightward asymmetry) of FA in cingulum (CING), uncinate (UNC), anterior limb of internal capsule (ALIC), external capsule (EC), posterior thalamic radiation/optic tract (PTR/OR), superior fronto-occipital fasciculus (SFOF), anterior and superior corona radiata (ACR and SCR), and the statistical significance with one-sample t test and effect sizes.

**Figure 6.** Left-right asymmetries of white matter (WM) tract fractional anisotropy (FA) values in infants and in 5-year-olds, measured from left-right tract-based spatial statistics (TBSS) skeleton by extracting with JHU-ICBM Atlas. Subpopulation of 22 subjects with MR scanning both at 2-5 weeks after birth and at age of 5 years. Positive values denote for leftward and negative values for rightward asymmetry. UNC = uncinate, SFOF = superior fronto-occipital fasciculus, SLF = superior longitudinal fasciculus, ST = stria terminalis, CING = cingulum, CG = cingulate, EC = external capsule, ILF/IFOF = inferior longitudinal fasciculus/inferior fronto-occipital fasciculus, PTR (OR) = posterior thalamic radiation (optic tract), PCR = posterior corona radiata, SCR = superior corona radiata, ACR = anterior corona radiata, rl IC = retrolenticular internal capsule, PLIC = posterior limb of internal capsule, ALIC = anterior limb of internal capsule, sup = superior, inf = inferior, cereb = cerebellar, CST = corticospinal tract, SCC = splenium of corpus callosum, BCC = body of corpus callosum, GCC = genu of corpus callosum, PCT = pontine crossing tract, MCP = middle cerebellar peduncle.

Differences in left-right lateralization of WM tracts between infants and 5-year-olds was compared in subset of 22 subjects included in both age groups (Figure 6). The detected asymmetries were more distinct in 5-year-olds, and the direction of asymmetry did not remain similar in all tracts from birth to the age of 5 years. Leftward asymmetries were detected in both age groups in superior longitudinal fasciculus (SLF), cingulum (CING), and cingulate (CG) and rightward asymmetry, for example, in the ALIC. However, the initial rightward asymmetry of posterior limb of internal capsule (PLIC), corticospinal tracts (CST), and inferior longitudinal fasciculus/inferior fronto-occipital fasciculus (ILF/IFOF), was revealed to shift to leftward laterality. A similar pattern was detected when repeating the comparison with entire subpopulations of infants (N = 166) and 5-year-olds (N = 144). In the group of 22 that were included in both the infant cohort and the 5-year-olds, the difference between lateralized FA in infants and 5-year-olds were statistically significant (p < 0.002) for CST, PLIC, anterior corona radiata and CG. Mean FA values of each tract in infants and in 5-year-olds are provided in the Supplement Table 5.

## 4 Discussion

We found widespread sex differences in white matter FA values in our sample of healthy, typically developing ca. 5-year-olds. Females had higher FA in all areas with significant differences, which were widely scattered throughout WM tracts. Sex differences were not detected in infants. Additionally, significant hemispheric asymmetries of FA values were observed in multiple WM tracts both in infants and at the age of 5 years. Hemispheric lateralization pattern was detected not to stay stable from birth to the age of 5 years. Lastly, age was positively associated with FA values during 6^th^ year of life in mid-posterior part of the corpus callosum.

Sex differences in WM tract structure were expected based on prior studies (Table 1). The previous literature on sex difference during development is, however, inconclusive, as the age ranges, methods and results vary widely between studies. In part of prior studies, no sex differences were observed ^14,51,52^, and in longitudinal developmental studies the observed sex differences have argued to be minor^19,26^. In the current study, the differences were significantly wider than previously described, and higher FA in females was observed in all regions with differences. The result highlights the sexual dimorphism in brain structure during development with significant detectable differences in multiple regions at the age of five years. Instead of tract-specific differences detected in a part of previous studies, we detected FA values to be higher in females across all brain regions. One possible explanation for varying results in prior literature may be the transient dynamics of sex differences. In studies reporting higher FA values in specified tracts in males ^21,32,46^ the age ranges are also higher (from 8 years onwards). The WM development of males has been shown to proceed later but also with steeper rates achieving and, in part of tracts, outpacing females^10^. A large cohort of 9- to 65-year-old subjects also showed sex differences to diminish with age^90^, further explaining why differences might not have been observed in older samples. Widespread sex differences observed in our 5-year-old population were emphasized to the posterior and temporal parts of the brain and were statistically more significant in the right hemisphere. These results agree with the prior studies showing both earlier brain development in females ^18,32,42–47^ and developmental pattern of maturation progressing in posterior-to-anterior direction. Our result of higher FA in females in widespread regions provides new insight into the timing of sex-dependent WM development trajectories and implies that at the age of 5 years the WM integrity is higher in females.

Hemispheric asymmetries of WM microstructure appear early ^58^ and are also suggested to remain stable throughout the brain development ^30,91^. However, asymmetries in WM of healthy, normally-developing children are not inspected carefully across whole WM, even though alterations of asymmetry is associated with different conditions such as dyslexia^59^. Knowledge of normal variation in asymmetric WM features is an important prerequisite for making conclusions from alterations in asymmetry. One recent study by Stephens et al. has shown significant asymmetry in all WM regions inspected (arcuate, cingulum, uncinate, IFOF, ILF, SLF and SCC) during the first 6 years of life^30^. Other studies on WM asymmetries in pediatric populations have mainly focused on age groups over 5 years of age, the sample sizes have remained quite modest and mainly leftward laterality in limited WM tracts has been observed^19,29,51,92,93^. Our findings further consolidate the emergence of hemispheric asymmetries during WM development. Lateralization patterns were detected already during neonatal ages, but also changes in asymmetric WM structure were detected between birth and the age of 5 years. The pattern of 5-year-olds resembled considerably the WM asymmetry detected in adult population. We compared our results with two prior adult-population studies (Supplement Figure 9), a TBSS study by Takao et al.^91^ and a Fixel-Based Analysis (FBA) study by Honnedevasthana Arun et al. 2021^94^. However, according to our results, some alterations in asymmetric pattern seem to occur between neonatal period and the age of 5 years. In comparison to prior longitudinal study of 0 to 6-year-olds^30^, in which the lateralization was measured based on mean DTI indices of each tract, we also detected variation in asymmetry within the tracts. Mainly leftward lateralization of CG was repeated in our population, however, the asymmetries of certain WM tracts varied between infants and 5-year-olds suggesting further development of the lateralization pattern during first years of life. Similarly, as in prior adult-studies^57,94^, we did not find association between the lateralization pattern and sex. Taken together, these findings highlight the role of asymmetry in fundamental structure of WM and underline the importance of early childhood as an essential period in the appearance of diverging lateralization patterns that may contribute to variation in cognitive measures and executive functions^95^, language development^96^ and to predisposition to diseases such as ADHD^97^ and autism^74,98^.

Age from birth and gestational age were expectedly detected to correlate positively with FA values. Gestational age showed stronger correlation with FA after controlling for the age from birth, which is plausible given the extensive WM development and partial myelination already *in utero*. When inspected separately, age after birth showed significant positive association with FA in males but not in females. The result may partly stem from different sample sizes (89 males vs 77 females) and is addressed for future longitudinal studies with more optimal data for observing developmental changes. Also, significant positive correlation between the age and FA of BCC was detected even in a narrow age range between 5.1 and 5.8 years. The posterior BCC and the isthmus of CC contain the primary somatosensory and motor fibres99, and the finding of rapid increase in the integrity in this area at the age of five years may be related to, e.g. the concurrent development of motor skills and bimanual movements^34,100^. Previously, higher initial FA values of callosal fibres have been associated with lower increase rates after the age of two years compared to for example association tracts^10^. One explanatory factor for detecting distinct age-effect in our data specifically in the CC might be the highly uniform organization of callosal fibres rendering the area susceptible for finding group differences if they exist.

The current study has also some limitations. The sample size is acknowledged to be modest, and the results need to be repeated in larger population and in other ethnic groups. As there were only 22 subjects included in both subpopulations, and the MR scanner was updated between imaging, the study setting is not technically longitudinal, but the subpopulations are considered as separate groups.

## 5 Conclusions

Rapid increase of the WM integrity, i.e. increase in FA values, occurs during the first five years of life. At that developmental timing, broad sex differences in WM structure are observed: higher FA and thus presumably more advanced development is detected in multiple regions in females compared to males. Additionally, the WM structure is widely lateralized during early development, and at the age of five years the lateralization pattern resembles considerably the ones previously described in adult studies, which has clear differences to those observed in infants.

## Author contributions

VK: data collection (5-year-olds), data pre-processing, formal analyses, writing the manuscript draft. HM: designing data processing pipeline, writing the manuscript. EPP, AC, ES: data collection (5-year-olds), writing the manuscript. JDL and JS: designing the MR acquisition protocol, writing the manuscript. SJS: data collection (infants), writing the manuscript. RP: radiological evaluation of the MR images, writing the manuscript. TL: neurological evaluation of the infants, writing the manuscript. LK and HK: establishing the cohort, building infrastructure for carrying out the study, writing the manuscript. JJT: study conception, funding acquisition, supervision of VK, building the pre-processing pipelines, writing the manuscript.

## Declarations

### Ethics approval

The study was approved by the Ethics Committee of the Hospital District of Southwest Finland ((07.08.2018) §330, ETMK: 31/180/2011), and performed in accordance with the Declaration of Helsinki.

### Data availability

The Finnish law and ethical permissions do not allow open sharing of the data used in this study, but data access is possible via formal material transfer agreements (MTA). Investigators that wish to access the data are encouraged to contact Principal Investigator of the FinnBrain Birth Cohort study Hasse Karlsson.

### Informed consent

Both parents of the participating child signed a written informed consent form. The child assent was assured prior to neuroimaging visits.

## Supplementary material to

**Supplement Table 1.**
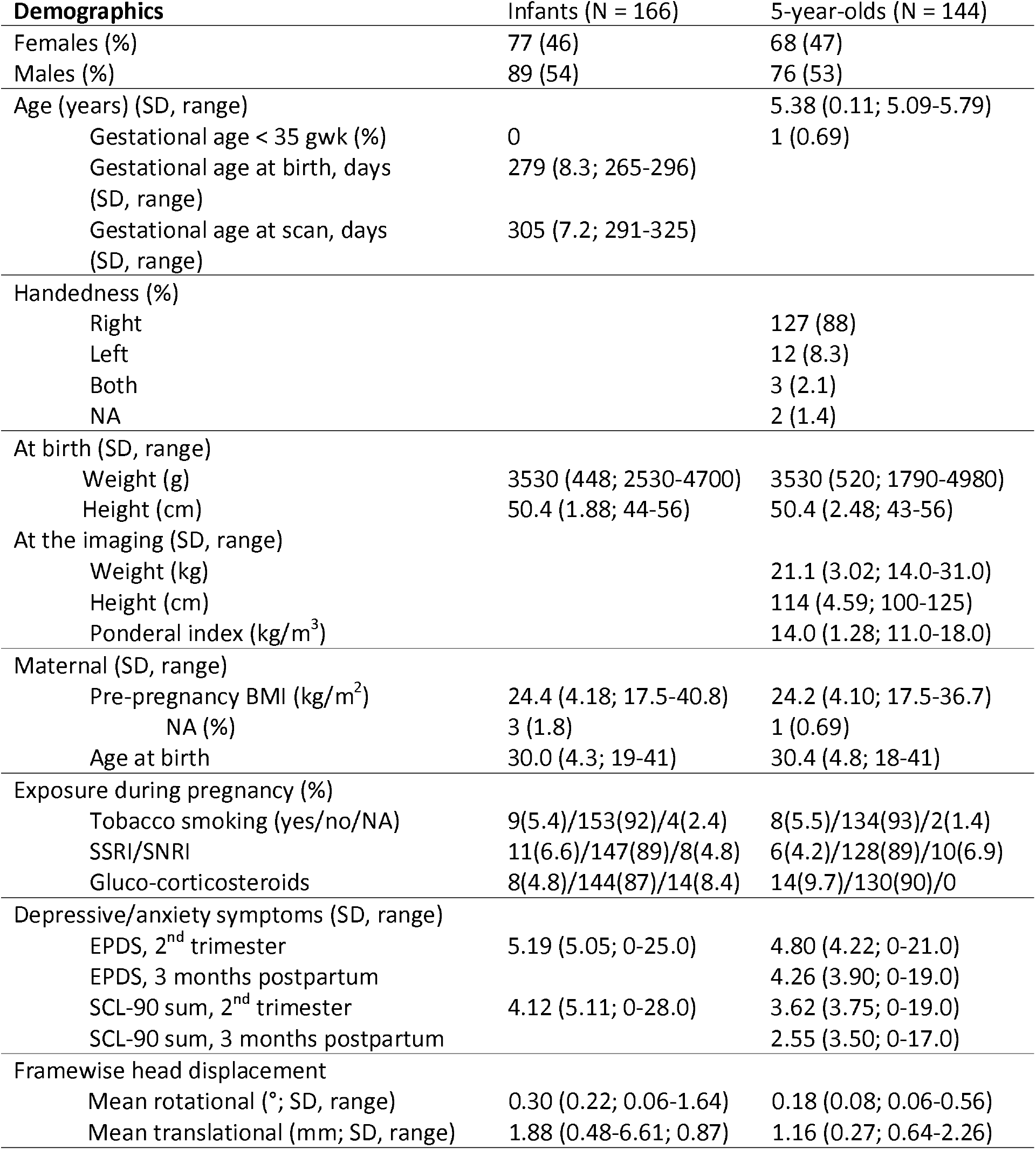
Demographics of the study population. SD = standard deviation, gwk = gestational week, BMI = body-mass index, SSRI/SNRI = selective serotonine/serotonine-noradrenaline re-uptake inhibitor, EPDS = Edinburgh postnatal depressive scale, SCL = symptom checklist, mm = millimetre

**Supplement Table 2.** Description of regression and sensitivity analyses in the current study. BMI = body-mass index, SES = socio-economic status, SSRI/SNRI = selective serotonine/serotonine-noradrenaline re-uptake inhibitor, EPDS = Edinburgh postnatal depressive scale, SCL = symptom checklist

| Regression analyses | Main variables | Covariates used in sensitivity analyses |
| --- | --- | --- |
| 5-year-olds | • sex<br>• age<br>• handedness<br>• Ponderal index<br>• Maternal pre-pregnancy BMI<br>• Maternal sosio-economis status (by educational status, categorised | • Gestational age<br>• Maternal age at birth<br>• Maternal pre-pregnancy BMI<br>• Maternal SES<br>• Smoking during pregnancy<br>• Exposure to glucocorticosteroids during pregnancy |
|  | high/medium to low)<br>Smoking during pregnancy<br>Intrascanner head motion | Exposure to SSRI/SNRI during pregnancy<br>EPDS at 2<sup>nd</sup> trimester<br>EPDS 3 months postpartum<br>SCL-90 at 2<sup>nd</sup> trimester<br>SCL-90 3 months postpartum |
| Infants | Age from birth<br>Gestational age<br>Sex<br>Birth weight<br>Maternal pre-pregnancy BMI<br>Maternal socio-economic status<br>Smoking during pregnancy<br>Intrascanner head motion | Gestational age<br>Age from birth<br>Maternal age at birth<br>Maternal pre-pregnancy BMI<br>Maternal SES<br>Smoking during pregnancy<br>Exposure to glucocorticosteroids during pregnancy<br>Exposure to SSRI/SNRI during pregnancy<br>EPDS at 2<sup>nd</sup> trimester<br>SCL-90 at 2<sup>nd</sup> trimester |

Example of sensitivity analyses

Example of sensitivity analyses
|  |  |
| --- | --- |
| Basic model | FA = age + sex + maternal BMI |
| Sensitivity model 1 | FA = age + sex + maternal BMI + maternal age |
| Sensitivity model 2 | FA = age + sex + maternal BMI + SES |
| Sensitivity model 3 | FA = age + sex + maternal BMI + smoking |
| Sensitivity model ... | FA = age + sex + maternal BMI + ... |

**Supplement Table 3.** Mean fractional anisotropy (FA) and standard deviation (SD) of each white matter tract and differences between girls and boys. CST = corticospinal tract, ML = medial lemniscus, ICP = inferior cerebellar peduncle, SCP = superior cerebellar peduncle, CP = cerebral peduncle, ALIC = anterior limb of internal capsule, PLIC = posterior limb of internal capsule, ACR = anterior corona radiata, SCR = superior corona radiata, PCR = posterior corona radiata, PTR (OR) = posterior thalamic radiation (optic tract), ILF/IFOF = inferior longitudinal fasciculus/inferior fronto-occipital fasciculus, EC = external capsule, CG = cingulate, CING = cingulum, ST = stria terminalis, SLF = superior longitudinal fasciculus, SFOF = superior fronto-occipital fasciculus, UNC= uncinate. Leftright difference of mean FA provided, negative values denote for rightward lateralization. Statistical significance calculated with independent sample t test (two-tailed), Bonferroni corrected p value = 0.001 (0.05/36), tracts with significant difference in bold.

|  |  | Mean<br>FA | SD | Sig. | Cohen's d |
| --- | --- | --- | --- | --- | --- |
| **Right** | **girl** | **0,587** | **0,029** | **0,000** | **0,680** |
| **PTR/OR** | **boy** | **0,566** | **0,032** |  |  |
| **Left** | **girl** | **0,587** | **0,030** | **0,000** | **0,658** |
| **PTR/OR** | **boy** | **0,566** | **0,034** |  |  |
| **Right** | **girl** | **0,516** | **0,028** | **0,000** | **0,644** |
| **ILF/IFOF** | **boy** | **0,499** | **0,026** |  |  |

|  |  | Mean |  |  |  |
| --- | --- | --- | --- | --- | --- |
|  |  | FA | SD | Sig. | Cohen's d |
| SCC | girl | 0,748 | 0,025 | 0,005 | 0,482 |
|  | boy | 0,735 | 0,029 |  |  |
| GCC | girl | 0,719 | 0,033 | 0,694 | 0,066 |
|  | boy | 0,721 | 0,036 |  |  |
| BCC | girl | 0,633 | 0,038 | 0,470 | 0,121 |
|  | boy | 0,637 | 0,037 |  |  |
| fornix | girl | 0,450 | 0,041 | 0,253 | 0,192 |
|  | boy | 0,442 | 0,043 |  |  |
| Right CST | girl | 0,488 | 0,024 | 0,666 | 0,072 |
|  | boy | 0,487 | 0,028 |  |  |
| Left CST | girl | 0,504 | 0,027 | 0,288 | 0,178 |
|  | boy | 0,509 | 0,028 |  |  |
| Right ALIC | girl | 0,544 | 0,023 | 0,725 | 0,059 |
|  | boy | 0,543 | 0,025 |  |  |
| Left ALIC | girl | 0,523 | 0,023 | 0,519 | 0,108 |
|  | boy | 0,520 | 0,027 |  |  |
| Right PLIC | girl | 0,654 | 0,020 | 0,570 | 0,095 |
|  | boy | 0,652 | 0,020 |  |  |
| Left PLIC | girl | 0,657 | 0,021 | 0,952 | 0,010 |
|  | boy | 0,657 | 0,021 |  |  |
| **Right rLIC** | **girl** | **0,553** | **0,028** | **0,001** | **0,582** |
|  | boy | 0,538 | 0,026 |  |  |
| Left rLIC | girl | 0,569 | 0,024 | 0,007 | 0,458 |
|  | boy | 0,559 | 0,023 |  |  |
| Right ACR | girl | 0,458 | 0,028 | 0,150 | 0,242 |
|  | boy | 0,452 | 0,028 |  |  |
| Left ACR | girl | 0,452 | 0,028 | 0,049 | 0,331 |
|  | boy | 0,443 | 0,027 |  |  |
| Right SCR | girl | 0,471 | 0,024 | 0,120 | 0,261 |
|  | boy | 0,464 | 0,025 |  |  |
| Left SCR | girl | 0,478 | 0,024 | 0,854 | 0,031 |
|  | boy | 0,477 | 0,022 |  |  |
| Right PCR | girl | 0,444 | 0,025 | 0,056 | 0,321 |
|  | boy | 0,436 | 0,028 |  |  |
| Left PCR | girl | 0,441 | 0,030 | 0,414 | 0,137 |
|  | boy | 0,437 | 0,027 |  |  |

| **Left** | **girl** | **0,517** | **0,026** | **0,001** | **0,579** |
| --- | --- | --- | --- | --- | --- |
| **ILF/IFOF** | **boy** | **0,502** | **0,026** |  |  |
| Right EC | girl | 0,381 | 0,020 | 0,311 | 0,170 |
|  | boy | 0,377 | 0,023 |  |  |
| Left EC | girl | 0,400 | 0,025 | 0,773 | 0,048 |
|  | boy | 0,401 | 0,020 |  |  |
| Right CING | girl | 0,458 | 0,035 | 0,463 | 0,123 |
|  | boy | 0,462 | 0,038 |  |  |
| Left CING | girl | 0,483 | 0,035 | 0,453 | 0,126 |
|  | boy | 0,488 | 0,039 |  |  |
| Right CG | girl | 0,409 | 0,039 | 0,860 | 0,029 |
|  | boy | 0,408 | 0,037 |  |  |
| Left CG | girl | 0,406 | 0,036 | 0,269 | 0,185 |
|  | boy | 0,412 | 0,037 |  |  |
| Right Fornix/ST | girl | 0,506 | 0,026 | 0,851 | 0,031 |
|  | boy | 0,505 | 0,032 |  |  |
| Left Fornix/ST | girl | 0,521 | 0,025 | 0,955 | 0,009 |
|  | boy | 0,521 | 0,032 |  |  |
| Right SLF | girl | 0,476 | 0,029 | 0,014 | 0,416 |
|  | boy | 0,465 | 0,025 |  |  |
| Left SLF | girl | 0,477 | 0,029 | 0,097 | 0,279 |
|  | boy | 0,469 | 0,025 |  |  |
| Right SFOF | girl | 0,478 | 0,036 | 0,469 | 0,121 |
|  | boy | 0,473 | 0,041 |  |  |
| Left SFOF | girl | 0,446 | 0,037 | 0,709 | 0,062 |
|  | boy | 0,449 | 0,045 |  |  |
| Right UNC | girl | 0,445 | 0,030 | 0,031 | 0,364 |
|  | boy | 0,434 | 0,028 |  |  |
| Left UNC | girl | 0,453 | 0,035 | 0,194 | 0,218 |
|  | boy | 0,447 | 0,028 |  |  |

**Supplement Figure 1.**
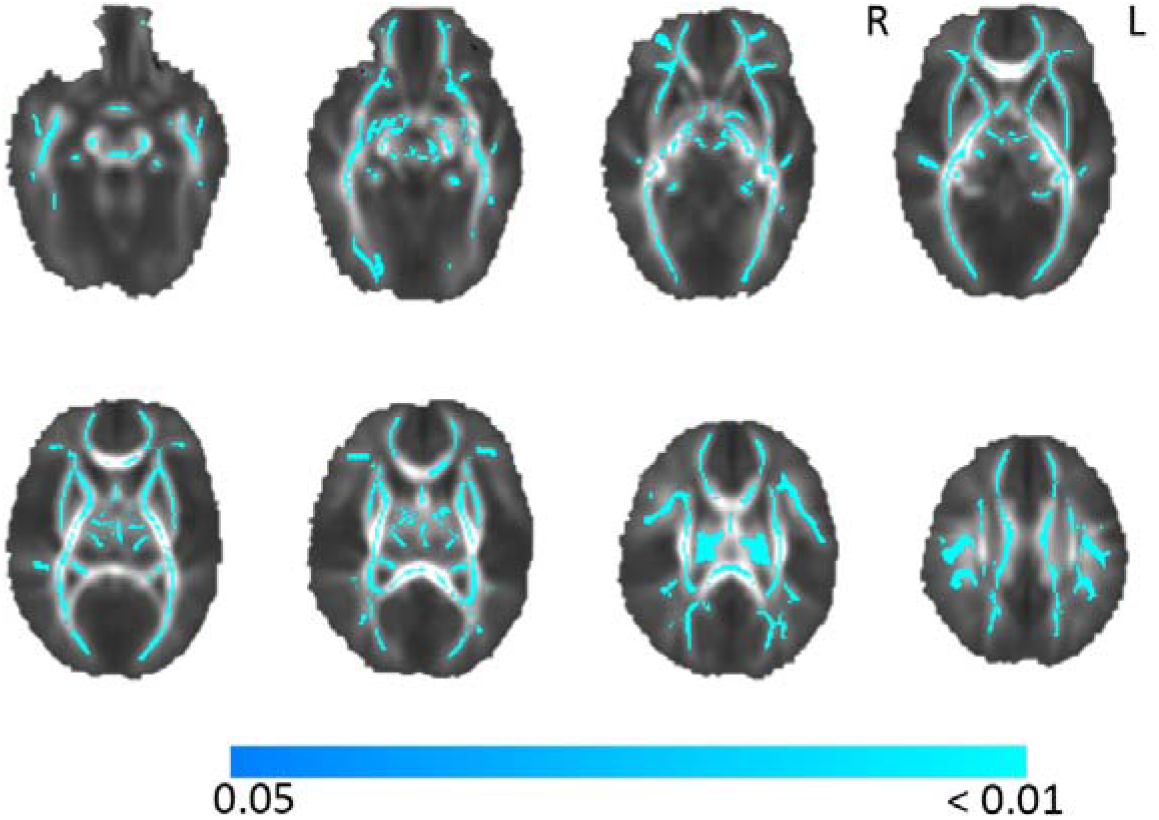
Effect of gestational age on fractional anisotropy analysed voxel-wise with tract-based spatial statistics with covariates (age after birth, sex, birth weight, intrascanner head motion, maternal tobacco smoking during pregnancy, socioeconomic status (low-medium/high), maternal prepregnancy body-mass index. Analysis conducted with 5000 permutations, threshold-free cluster enhancement correction applied, p < 0.05, R = right, L = left.

**Supplement Figure 2.**
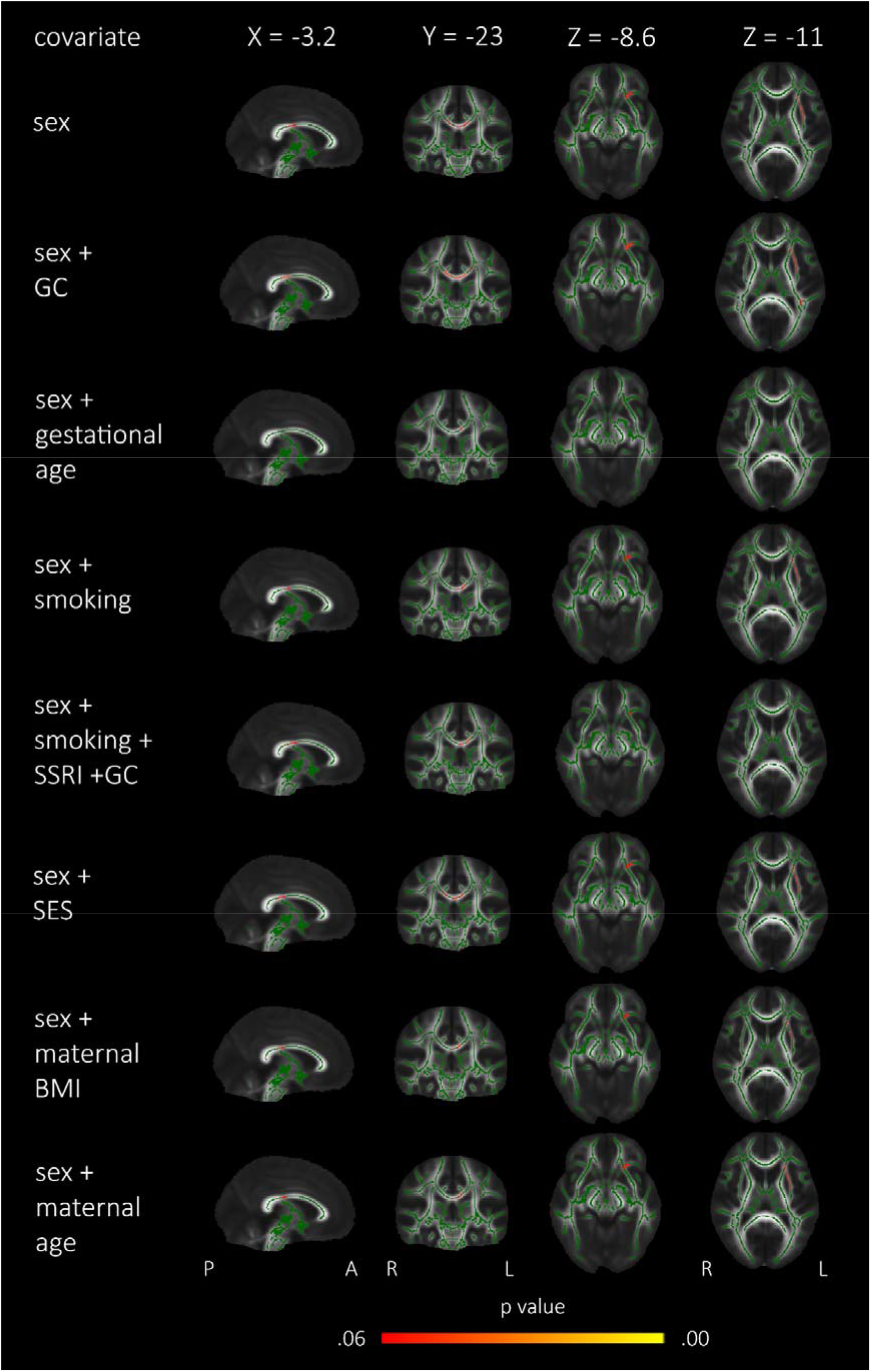
Age-effects analysed voxel-wise with tract-based spatial statistics with covariates (prenatal glucocorticoid (GC) exposure, gestational age, maternal tobacco smoking during pregnancy, exposure to selective serotonin reuptake inhibitors (SSRI) during pregnancy, socioeconomic status (SES; low-medium/high), maternal pre-pregnancy bodymass index (BMI) and maternal age at birth). Analyses conducted with 5000 permutations, TFCE (threshold-free cluster enhancement) correction applied, p < 0.06.

**Supplement Figure 3.**
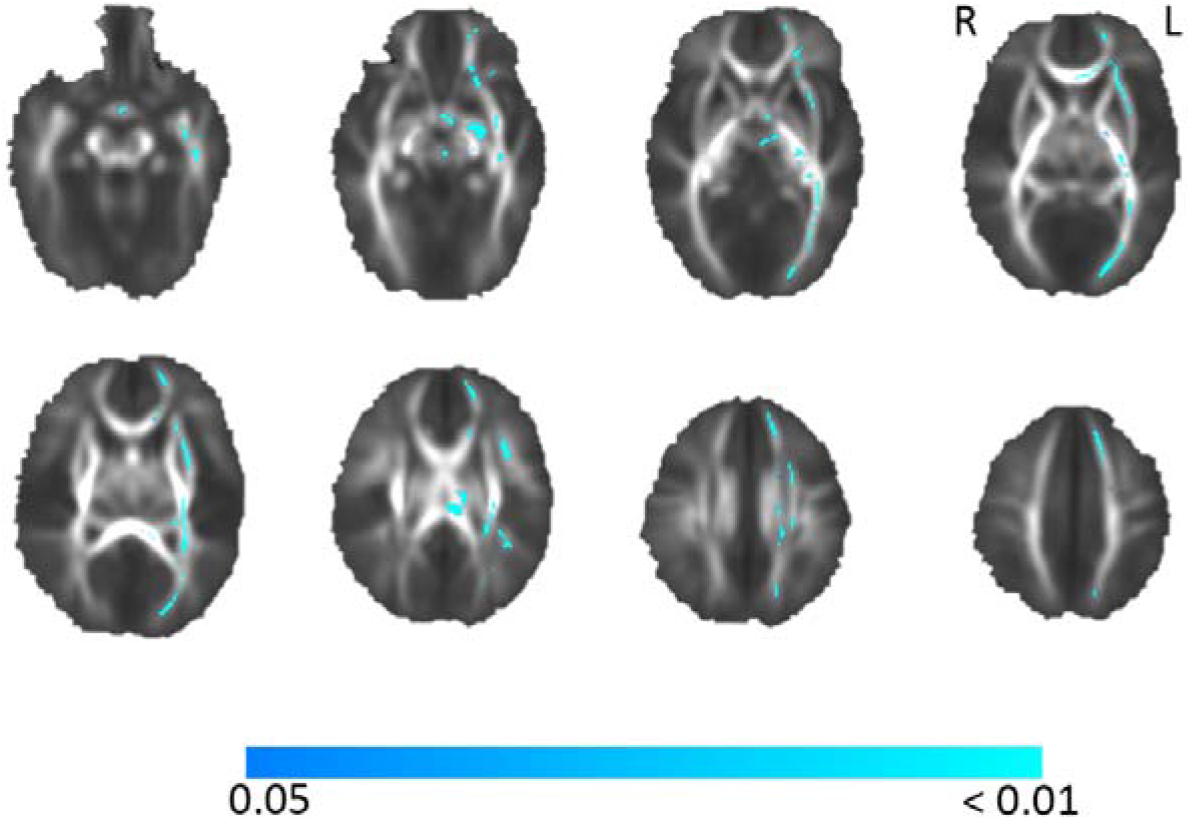
Lateralization pattern of white matter tract fractional anisotropy with tract-based spatial statistics in infants. Significant regions with leftward asymmetry in blue. Threshold-free cluster enhancement (TFCE) correction applied, 5000 permutations, p < 0.05 (blue colour bar showing p value). R = right, L = left

**Supplement Figure 4.**
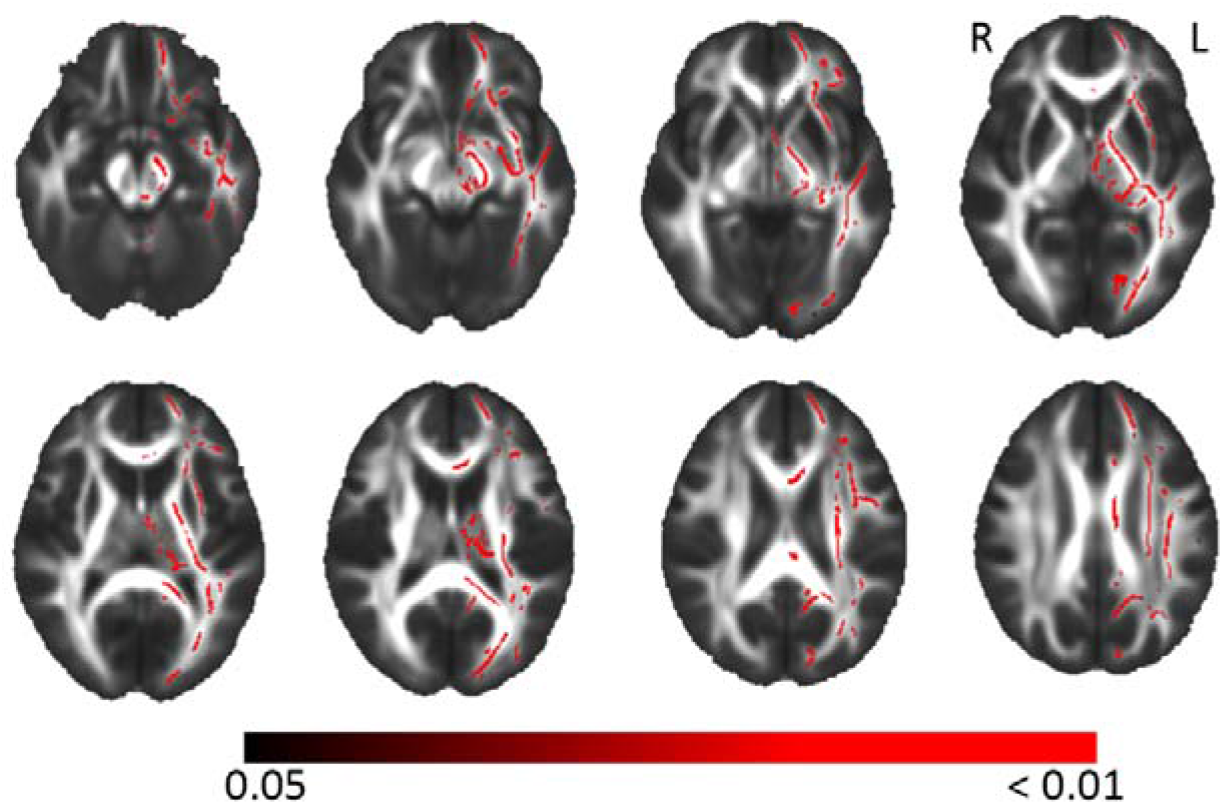
Lateralization pattern of white matter tract fractional anisotropy with tract-based spatial statistics in 5-year-olds. Significant regions with leftward asymmetry in red. Threshold-free cluster enhancement (TFCE) correction applied, 5000 permutations, p < 0.05 (red colour bar showing p value). R = right, L = left

**Supplement Figure 5.**
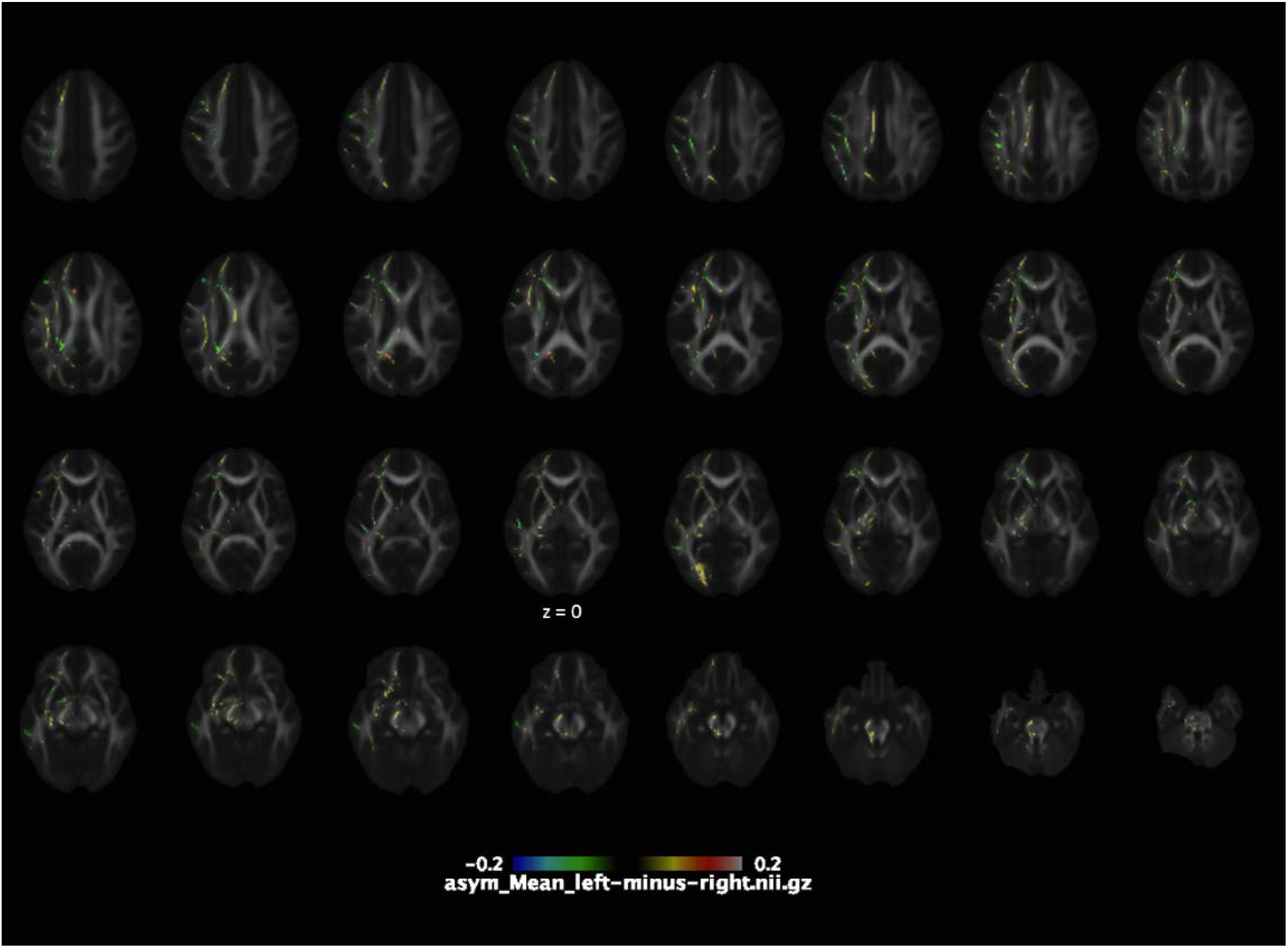
Mean values of lateralized FA (left minus right). Yellow to red colours (positive values) indicate leftward asymmetry and blue to green colours (negative values) rightward asymmetry.

**Supplement Figure 6.**
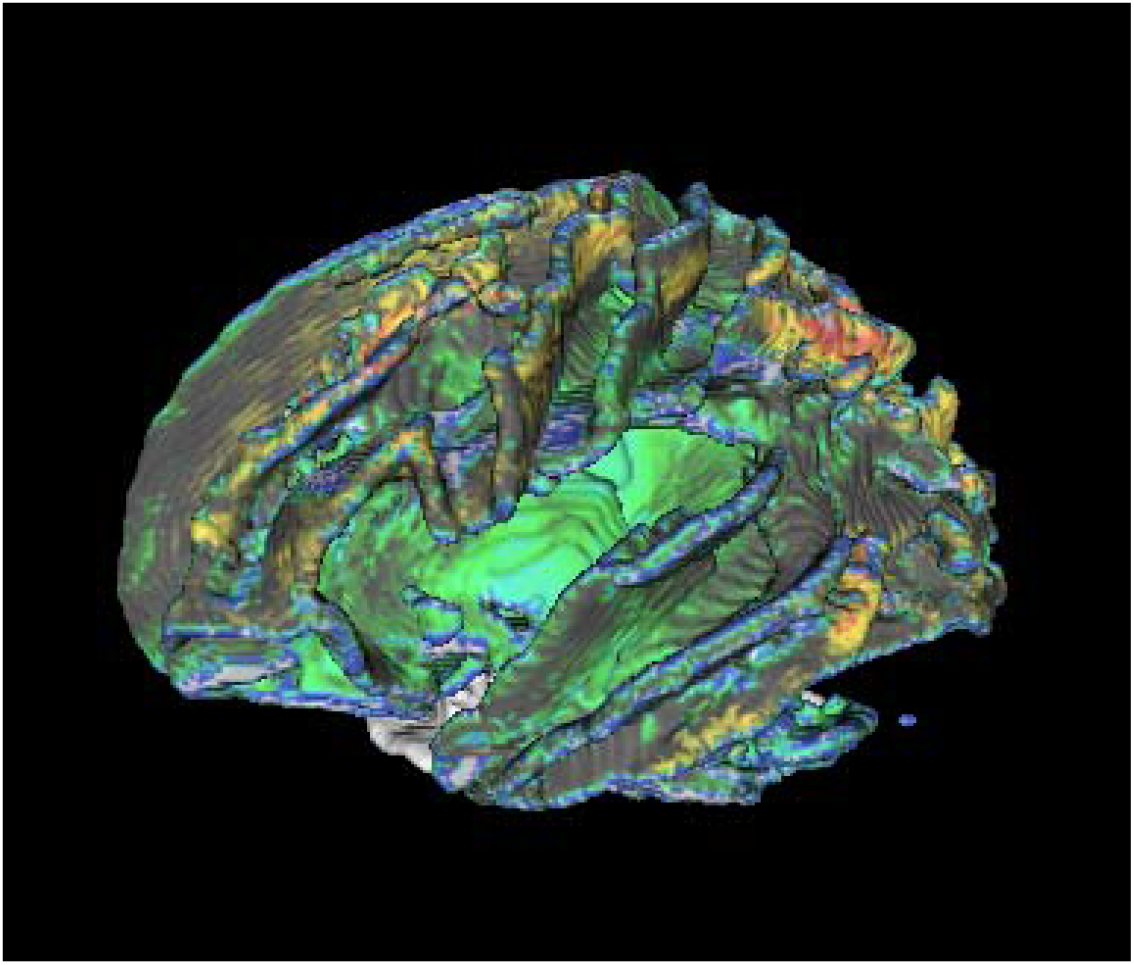
Yellow to red colours (positive values) indicate leftward asymmetry and blue to green colours (negative values) rightward asymmetry.

**Supplement Figure 7.**
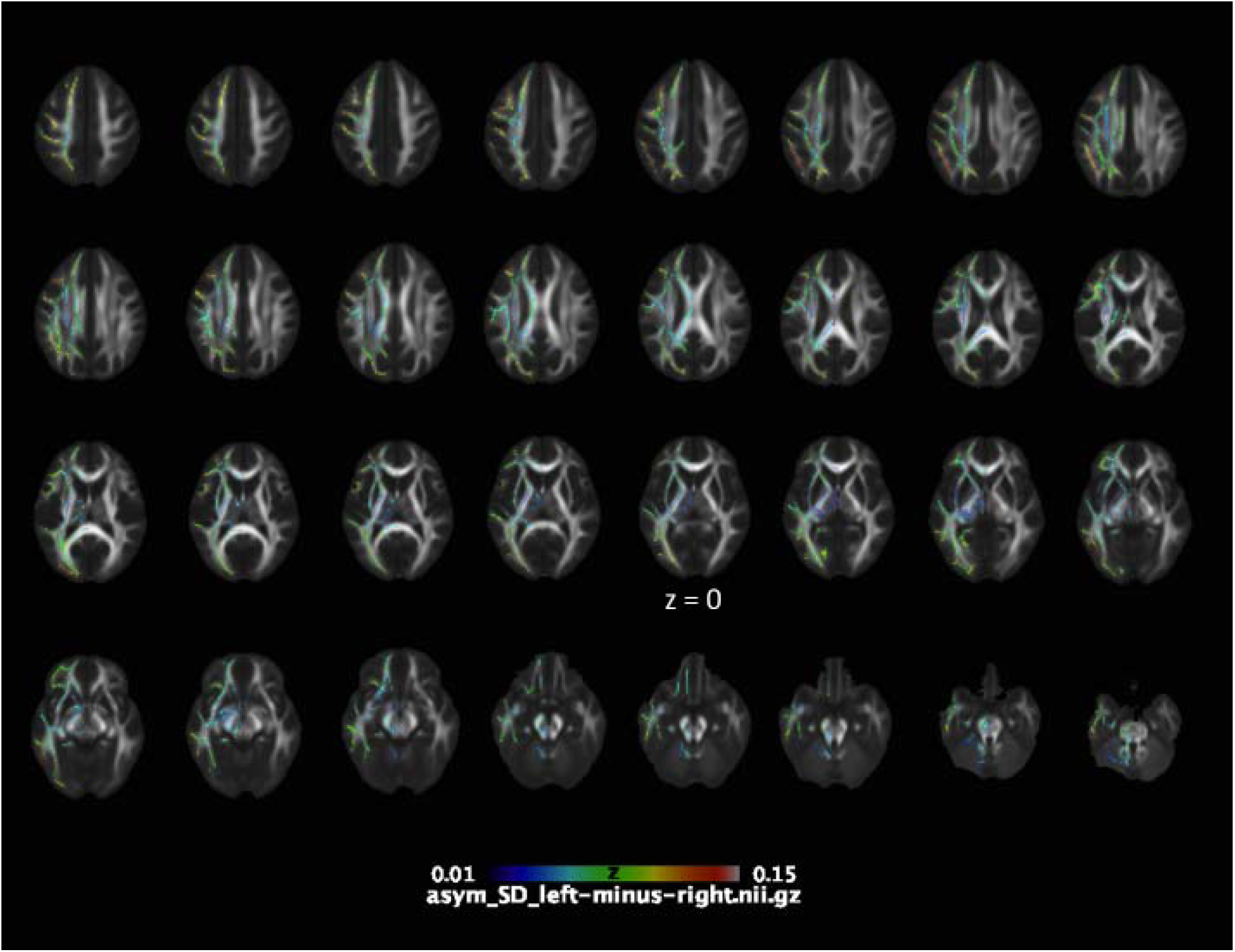
Inter-subject variation of lateralized FA values described as standard deviation (SD).

**Supplement Table 4.** Comparison of mean fractional anisotropy (FA) values in bilateral white matter tracts in 5-year-olds. CST = corticospinal tract, ML = medial lemniscus, ICP = inferior cerebellar peduncle, SCP = superior cerebellar peduncle, CP = cerebral peduncle, ALIC = anterior limb of internal capsule, PLIC = posterior limb of internal capsule, ACR = anterior corona radiata, SCR = superior corona radiata, PCR = posterior corona radiata, PTR (OR) = posterior thalamic radiation (optic tract), ILF/IFOF = inferior longitudinal fasciculus/inferior fronto-occipital fasciculus, EC = external capsule, CG = cingulate, CING = cingulum, ST = stria terminalis, SLF = superior longitudinal fasciculus, SFOF = superior fronto-occipital fasciculus, UNC = uncinate. Left-right difference of mean FA provided, negative values denote for rightward lateralization. Statistical significance calculated with one sample t test (two-tailed), Bonferroni corrected p value = 0.001 (0.05/36), tracts with significant difference in bold.

| Tract |  | Mean FA | SD | Left-right difference | Sig. | Cohen's d |
| --- | --- | --- | --- | --- | --- | --- |
| **CST** | right | **0,487** | **0,026** | **0,0194** | **0,000** | **0,721** |
|  | left | **0,507** | **0,028** |  |  |  |
| **ALIC** | right | **0,544** | **0,024** | **-0,0224** | **0,000** | **0,903** |
|  | left | **0,521** | **0,025** |  |  |  |
| PLIC | right | 0,653 | 0,020 | 0,0041 | 0,087 | 0,202 |
|  | left | 0,657 | 0,021 |  |  |  |
| ACR | right | 0,455 | 0,028 | -0,0077 | 0,019 | 0,278 |
|  | left | 0,447 | 0,028 |  |  |  |
| **SCR** | right | **0,467** | **0,025** | **0,0100** | **0,000** | **0,420** |
|  | left | 0,477 | 0,023 |  |  |  |
| PCR | right | 0,440 | 0,027 | -0,0006 | 0,843 | 0,023 |
|  | left | 0,439 | 0,028 |  |  |  |
| PTR/OR | right | 0,576 | 0,032 | -0,0002 | 0,951 | 0,007 |
|  | left | 0,576 | 0,034 |  |  |  |
| ILF/IFOF | right | 0,507 | 0,028 | 0,0021 | 0,515 | 0,077 |
|  | left | 0,509 | 0,027 |  |  |  |
| **EC** | right | **0,379** | **0,022** | **0,0220** | **0,000** | **1,00** |
|  | left | 0,401 | 0,022 |  |  |  |
| **CING** | right | **0,460** | **0,036** | **0,0256** | **0,000** | **0,700** |
|  | left | 0,485 | 0,037 |  |  |  |
| CG | right | 0,408 | 0,038 | 0,0010 | 0,822 | 0,026 |
|  | left | 0,409 | 0,036 |  |  |  |
| SLF | right | 0,470 | 0,027 | 0,0027 | 0,403 | 0,099 |
|  | left | 0,473 | 0,027 |  |  |  |
| **SFOF** | right | **0,475** | **0,039** | **-0,0281** | **0,000** | **0,706** |
|  | left | **0,447** | **0,041** |  |  |  |
| UNC | right | 0,439 | 0,029 | 0,0106 | 0,003 | 0,350 |
|  | left | 0,450 | 0,031 |  |  |  |

**Supplement Figure 8.**
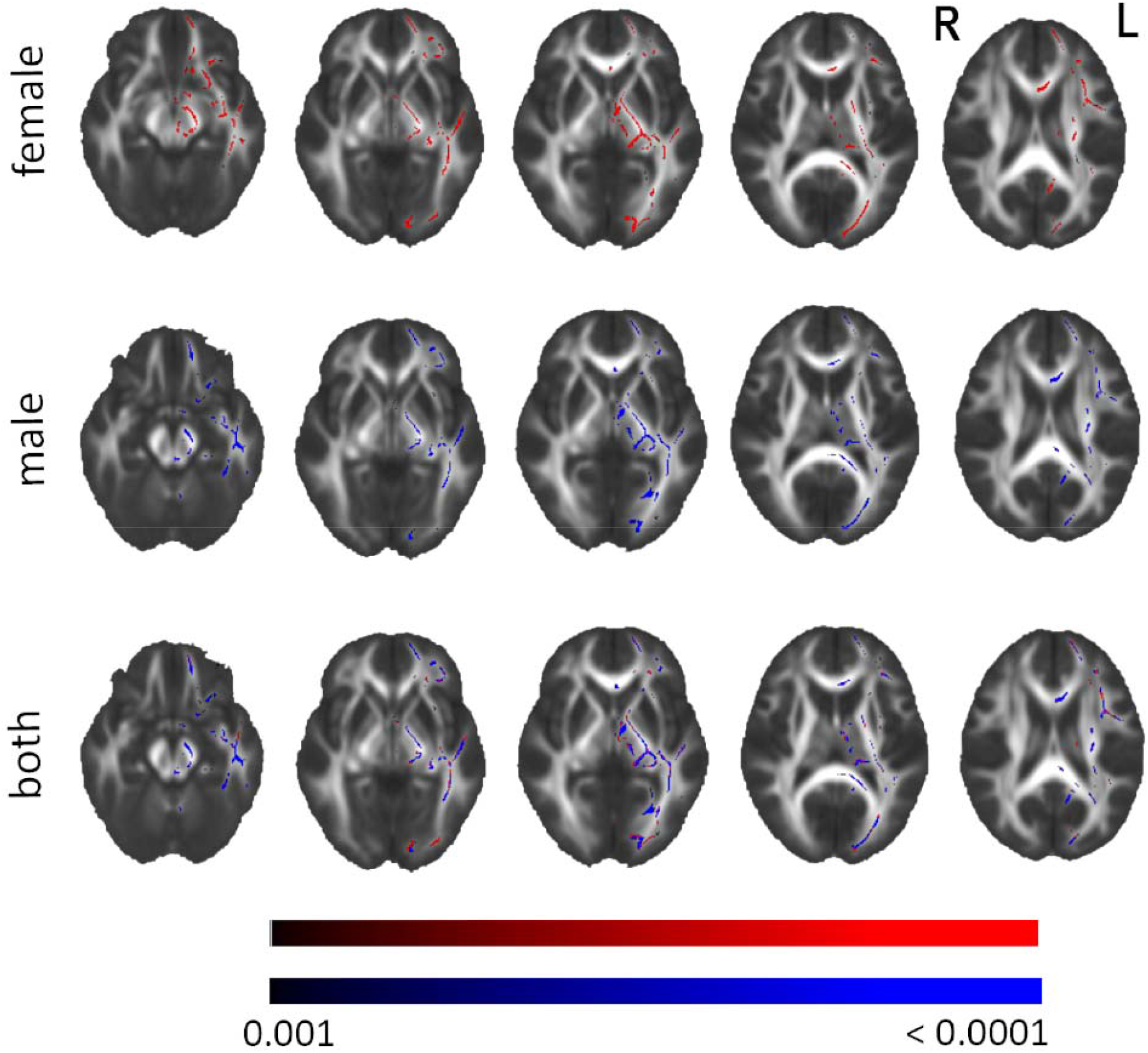
Sex-specific white matter lateralization patterns of 5-year-olds with Tract-based Spatial Statistical (TBSS) analysis, depicted as left-right fractional anisotropy values. The statistically significantly asymmetric regions showed with red in females and with blue in males. Analyses conducted with 5000 permutations, threshold-free cluster enhancement (TFCE) correction applied, p < 0.001. R = right, L = left

**Supplement Table 5.**
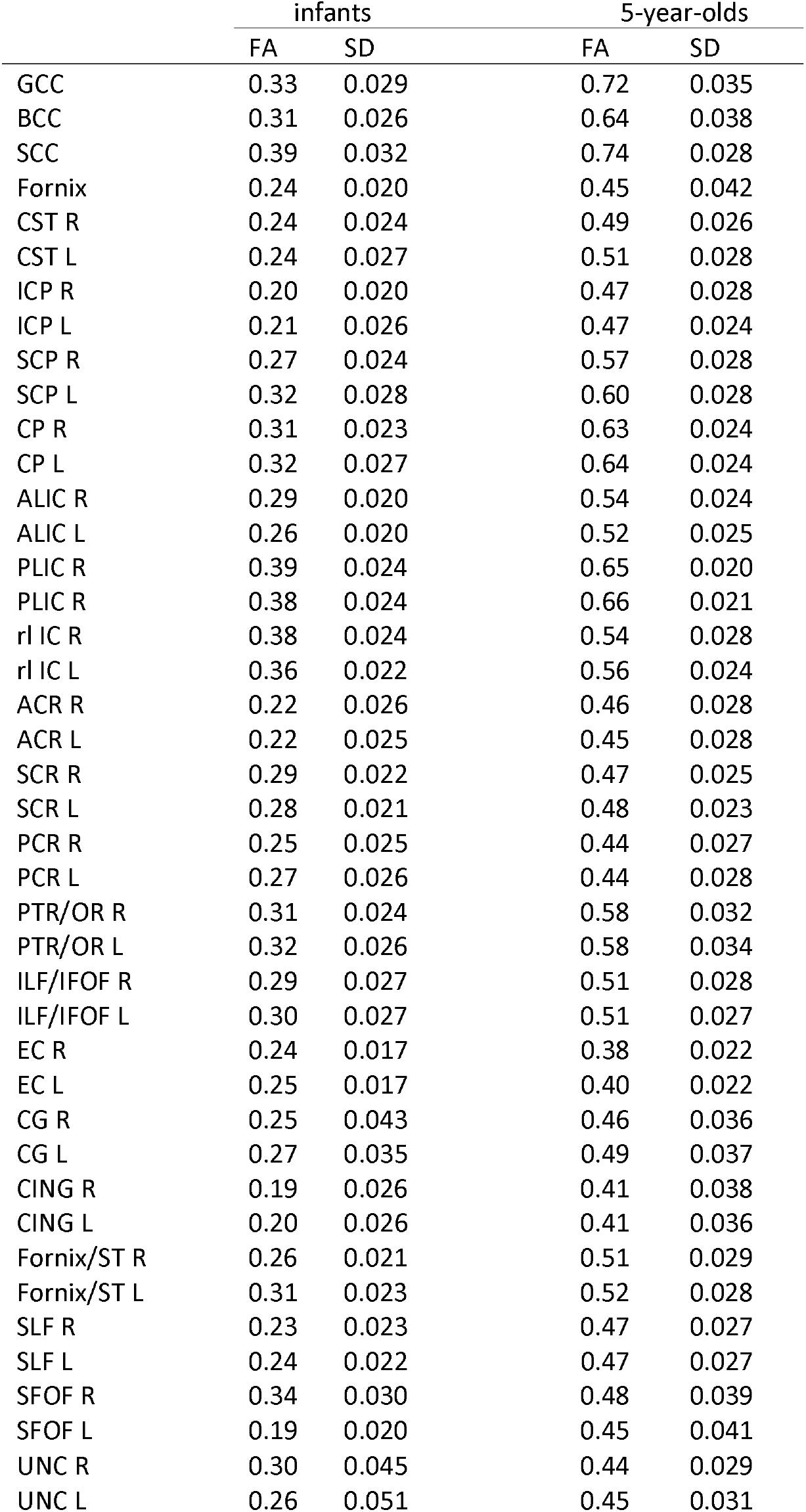
Mean fractional anisotropy (FA) and standard deviation (SD) of each white matter tract in infants and in 5-year-olds. GCC = genu of corpus callosum, BCC = body of corpus callosum, SCC = splenium of corpus callosum, CST = corticospinal tract, ICP = inferior cerebellar peduncle, SCP = superior cerebellar peduncle, CP = cerebral peduncle, ALIC = anterior limb of internal capsule, PLIC = posterior limb of internal capsule, rl IC = retrolenticular internal capsule, ACR = anterior corona radiata, SCR = superior corona radiata, PCR = posterior corona radiata, PTR (OR) = posterior thalamic radiation (optic tract), ILF/IFOF = inferior longitudinal fasciculus/inferior fronto-occipital fasciculus, EC = external capsule, CG = cingulate, CING = cingulum, ST = stria terminalis, SLF = superior longitudinal fasciculus, SFOF = superior fronto-occipital fasciculus, UNC = uncinate, R = right, L = left.

|  | infants |  | 5-year-olds |  |
| --- | --- | --- | --- | --- |
|  | FA | SD | FA | SD |
| GCC | 0.33 | 0.029 | 0.72 | 0.035 |
| BCC | 0.31 | 0.026 | 0.64 | 0.038 |
| SCC | 0.39 | 0.032 | 0.74 | 0.028 |
| Fornix | 0.24 | 0.020 | 0.45 | 0.042 |
| CST R | 0.24 | 0.024 | 0.49 | 0.026 |
| CST L | 0.24 | 0.027 | 0.51 | 0.028 |
| ICP R | 0.20 | 0.020 | 0.47 | 0.028 |
| ICP L | 0.21 | 0.026 | 0.47 | 0.024 |
| SCP R | 0.27 | 0.024 | 0.57 | 0.028 |
| SCP L | 0.32 | 0.028 | 0.60 | 0.028 |
| CP R | 0.31 | 0.023 | 0.63 | 0.024 |
| CP L | 0.32 | 0.027 | 0.64 | 0.024 |
| ALIC R | 0.29 | 0.020 | 0.54 | 0.024 |
| ALIC L | 0.26 | 0.020 | 0.52 | 0.025 |
| PLIC R | 0.39 | 0.024 | 0.65 | 0.020 |
| PLIC L | 0.38 | 0.024 | 0.66 | 0.021 |
| rl IC R | 0.38 | 0.024 | 0.54 | 0.028 |
| rl IC L | 0.36 | 0.022 | 0.56 | 0.024 |
| ACR R | 0.22 | 0.026 | 0.46 | 0.028 |
| ACR L | 0.22 | 0.025 | 0.45 | 0.028 |
| SCR R | 0.29 | 0.022 | 0.47 | 0.025 |
| SCR L | 0.28 | 0.021 | 0.48 | 0.023 |
| PCR R | 0.25 | 0.025 | 0.44 | 0.027 |
| PCR L | 0.27 | 0.026 | 0.44 | 0.028 |
| PTR/OR R | 0.31 | 0.024 | 0.58 | 0.032 |
| PTR/OR L | 0.32 | 0.026 | 0.58 | 0.034 |
| ILF/IFOF R | 0.29 | 0.027 | 0.51 | 0.028 |
| ILF/IFOF L | 0.30 | 0.027 | 0.51 | 0.027 |
| EC R | 0.24 | 0.017 | 0.38 | 0.022 |
| EC L | 0.25 | 0.017 | 0.40 | 0.022 |
| CG R | 0.25 | 0.043 | 0.46 | 0.036 |
| CG L | 0.27 | 0.035 | 0.49 | 0.037 |
| CING R | 0.19 | 0.026 | 0.41 | 0.038 |
| CING L | 0.20 | 0.026 | 0.41 | 0.036 |
| Fornix/ST R | 0.26 | 0.021 | 0.51 | 0.029 |
| Fornix/ST L | 0.31 | 0.023 | 0.52 | 0.028 |
| SLF R | 0.23 | 0.023 | 0.47 | 0.027 |
| SLF L | 0.24 | 0.022 | 0.47 | 0.027 |
| SFOF R | 0.34 | 0.030 | 0.48 | 0.039 |
| SFOF L | 0.19 | 0.020 | 0.45 | 0.041 |
| UNC R | 0.30 | 0.045 | 0.44 | 0.029 |
| UNC L | 0.26 | 0.051 | 0.45 | 0.031 |

**Supplement Figure 9.**
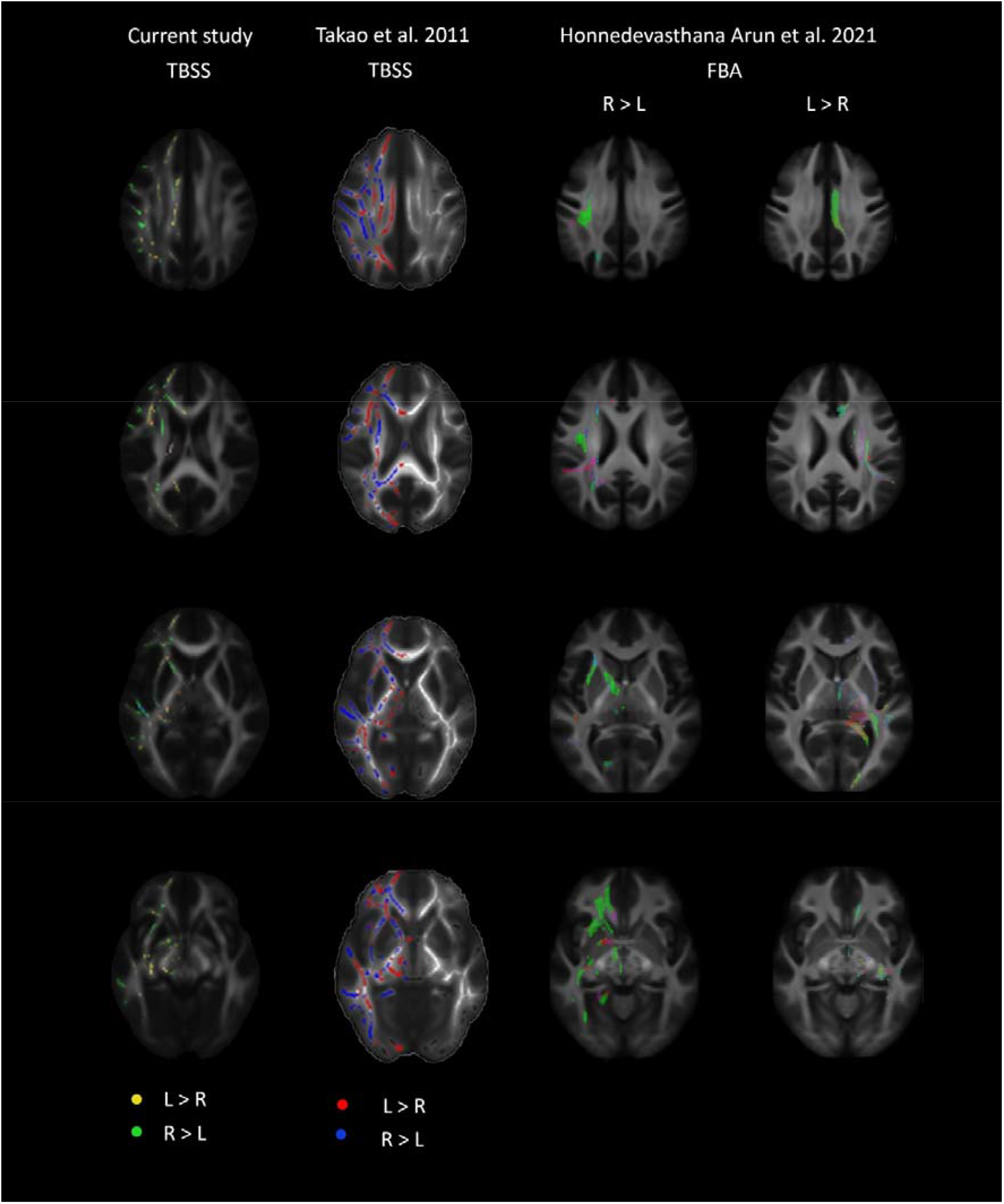
Comparison of lateralization patterns in our study and two previously published adult-population studies (Takao et al. 2011 and Honnedevasthana Arun et al. 2021). Figure adapted from previously published figures by Takao et al. 2011^1^ and Honnedevasthana Arun et al. 2021^2^, Copyright Elsevier. TBSS = tract-based spatial statistics, FBA = fixel-based analysis, L = left, R = right

